# Anterior Cleft Palate due to Cbfb deficiency and its rescue by folic acid

**DOI:** 10.1101/515650

**Authors:** Safiye E. Sarper, Toshihiro Inubushi, Hiroshi Kurosaka, Hitomi Ono Minagi, Yuka Murata, Koh-ichi Kuremoto, Takayoshi Sakai, Ichiro Taniuchi, Takashi Yamashiro

## Abstract

Core binding factor β (Cbfb) is a cofactor of Runx transcription factors. Among Runx transcription factors, Runx1 is a prerequisite for anterior-specific palatal fusion. However, whether Cbfb serves as a modulator or obligatory factor in Runx signaling that regulates palatogenesis is unclear. We herein report that Cbfb is essential and indispensable in anterior palatogenesis. Palatal fusion in *Cbfb* mutants is disturbed due to failed disintegration of the fusing epithelium specifically at the anterior portion, as is observed in *Runx1* mutants. In this mutants, the *Tgfb3* expression is disturbed at the corresponding area of the failed palatal fusion, where phosphorylation of Stat3 is also disturbed. TGFB3 protein rescues the palatal fusion *in vitro.* Strikingly, the anterior cleft palate in *Cbfb* mutants is further rescued by pharmaceutical application of folic acid that activates suppressed Stat3 phosphorylation and *Tgfb3* expression *in vitro.* With these findings, we provide the first evidence that Cbfb is a prerequisite for anterior palatogenesis as an obligatory cofactor in the Runx1/Cbfb-Stat3-Tgfb3 signaling axis. Furthermore, the rescue of the mutant cleft palate using folic acid may elucidate potential therapeutic targets by Stat3 modification for the prevention and pharmaceutical intervention of cleft palate.

**Summary Statement:** Epithelial deletion of Cbfb results in an anterior cleft palate with impaired fusion of the palatal process and folic acid application rescues the mutant phenotype with Stat3 activation *in vitro.*

## Introduction

Cleft palate is the most common congenital anomalies in humans, and its etiology is complex (Dixon et al., 2011; Murray, 2002). The palate is derived from the primary and secondary palate, which are located in the anterior and posterior portions of the palate, respectively (Gu et al., 2008). Palatal fusion is essential in palatogenesis, and its defect leads to cleft palate. Two halves of the palatal process fuse in the middle to form the secondary palate, which further fuse with the primary palate and the nasal septum to form the definite palate (Ferguson, 1988).

Among various molecules regulating palatogenesis, Runx1 is involved in the regulation of palatal fusion specifically in the anterior region. Epithelial-specific loss of *Runx1* results in failure of palatal fusion specifically at the anterior portion between the primary and the secondary palate with failed disintegration of the medial-edge epithelium. In this mutants, the *Tgfb3* expression was disturbed among various molecules regulating palatogenesis, and Stat3 phosphorylation was also downregulated (Sarper et al., 2018). Since TGFB3 protein rescues the mutant cleft palate, Tgfb3 is critical in Runx1 signaling in palatogenesis (Sarper et al., 2018). Furthermore, Stat3 inhibitor disturbed the palatal fusion accompanied by the downregulation on the *Tgfb3* expression, suggesting that extrinsic modification of the Stat3 activity affects Tgfb3 signaling and may be a potential therapeutic target in pharmaceutical intervention for cleft palate (Sarper et al., 2018).

Core binding factor β (Cbfb) is a cofactor of Runx family genes (*Runx1, Runx2* and *Runx3*) that forms a heterodimeric transcription complex (Huang et al., 2001). Cbfb enhances the binding affinity to DNA and also promotes Runx protein stability (Huang et al., 2001; Ogawa et al., 1993; Wang et al., 1993). Of note, Cbfb can act as either an obligate cofactor for the Runx function or a dispensable modulator of the Runx activity (Gau et al., 2017). For example, Cbfb acts as an obligate cofactor for the Runx function in hematopoietic cells (Chen et al., 2011) but as a dispensable modulator of the Runx activity in skeletogenesis (Yoshida et al., 2002). However, the possible functional role of Cbfb in palatogenesis has not been investigated.

A human genome study demonstrated that *CBFb* haploinsufficiency due to an interstitial deletion caused cleft palate and congenital heart anomalies in human (Khan et al., 2006; Tsoutsou et al., 2013; Yamamoto et al., 2008). A chromosomal fragile site of FRA16B, which co-localizes with breakpoints within *CBFb* at the chromosomal locus 16q22.1., is also involved in the inheritance of cleft palate (McKenzie et al., 2002). However, whether Cbfb is an obligate cofactor or a dispensable modulator in Runx1 signaling in palatogenesis has not been investigated.

Maternal folic acid supplementation has been shown to be as an effective intervention for reducing the risk of non-syndromic cleft palate (Millacura et al., 2017; Wehby and Murray, 2010). However, the mechanism by which folic acid prevents such structural anomalies in the fetus is still unknown (Obican et al., 2010). Interestingly, folic acid and folate can activate Stat3 (Hansen et al., 2015; Wei et al., 2017). Our previous study has shown that pharmaceutical application of Stat3 inhibitors disturbs the palatal fusion with downregulation of *Tgfb3.* Hence, it was assumed that folic acid might be a useful therapy for preventing the cleft palate via the extrinsic modification of Stat3 activation to prevent cleft palate.

We herein report the first evidence that Cbfb is essential in anterior palatogenesis as an obligatory cofactor in the Runx1/Cbfb-Stat3-Tgfb3 signaling axis. In addition, we also demonstrate the rescue of mutant cleft palate via pharmaceutical folic acid application, at least in part, by activating Stat3 phosphorylation in the Runx1/Cbfb-Tgfb3 signaling axis during palatogenesis.

## Results

### Palatal phenotypes in Cbfb mutants

The palatal phenotype was evaluated in vivo to see how Cbfb affect the palatal fusion using epithelial-specific conditional knockout mice (K14-Cre/Cbfb^fl/fl^) (Kurosaka et al., 2011).

The recombination efficiency of K14-Cre was evaluated in the developing palate previously (Sarper et al., 2018).

In this Cbfb mutants, anterior cleft was evident between the primary and secondary palates both at P0 and P50 (Fig. 1A-D). The cleft was seen in 100% of the mutants (n=8) when evaluated at P0 (Fig. 1E). In histological sections (Fig. 1F), failed palatal fusion was also confirmed at the first rugae A-P level in *Cbfb* mutants at E17.0 (Fig. 1G-J). In the more posterior portion, the secondary palate did not contact to the primary palate or the nasal septum (Fig. 1K,L). These findings show that the morphological palatal phenotypes are similar to those in *Runx1* mutants (Sarper et al., 2018).

**Figure 1.**
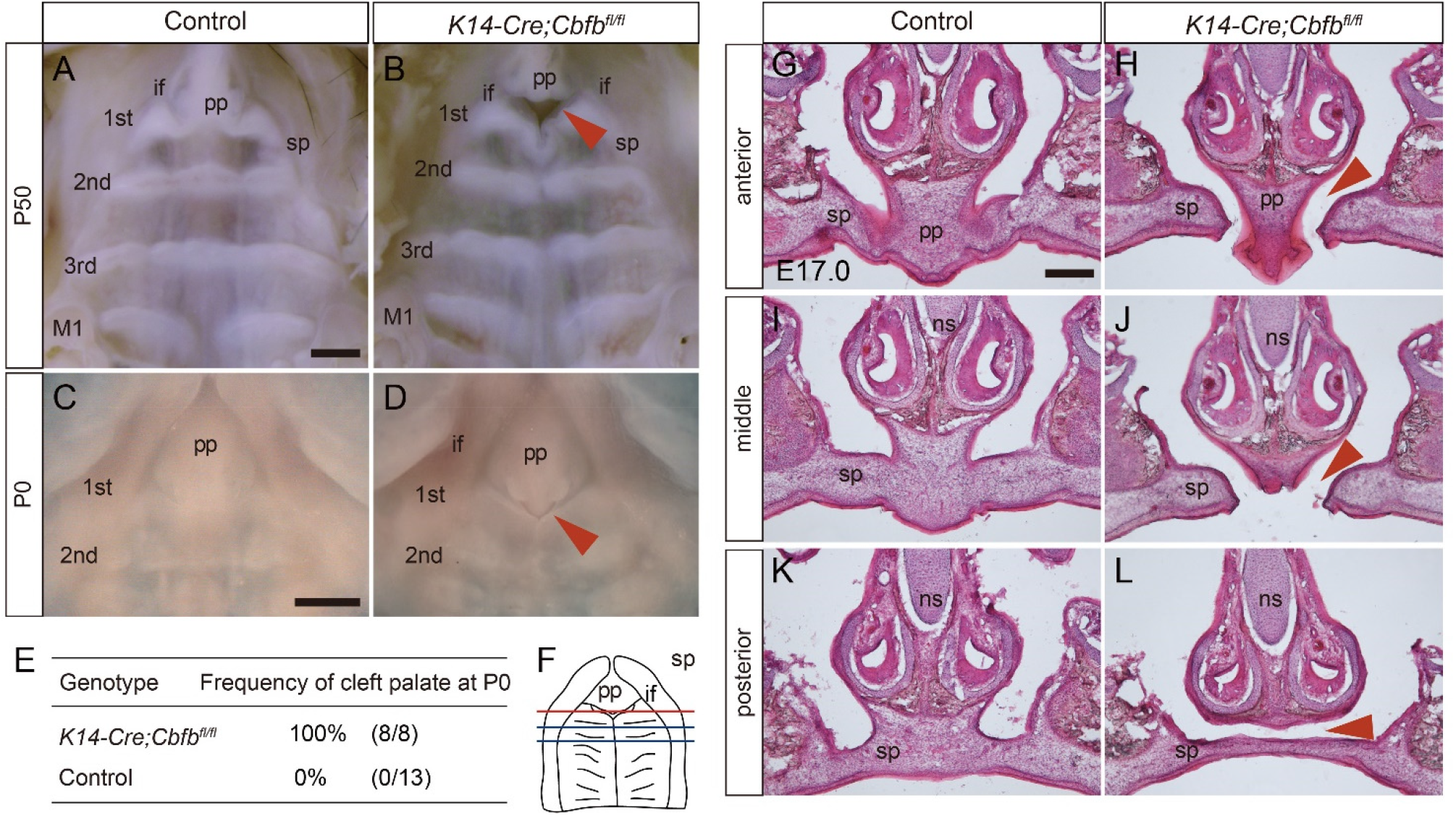
Palatal phenotypes of *K14-Cre/Cbfb^fl/fl^* mice. (A-D) Occlusal views of control and *Cbfb* mutant mouse palates. An anterior cleft palate was evident at the boundary between the primary and secondary palates in *Cbfb* mutant palates both at P50 (A,B) and P0 (C,D). The arrowheads indicate the cleft. Scale bar: 400 μm. (E) The table indicates the frequency of anterior cleft in control and *Cbfb* mutant mice at P0. (F) The diagram shows the occlusal view of the palate and the section positions as indicated by the lines. (G-L) Histological sections at E17.0 revealed that the palatal shelves of *Cbfb* mutant mice did not make contact at the boundary between the primary and secondary palate (G,H,J,K). In the more posterior region, the secondary palate was fused completely, however, the fused palate did not make contact with the inferior border of the nasal septum (I,L). Arrowheads indicate the failure of fusion. Scale bar: 200 μm.

### Characterization of the mutant epithelium in palatal fusion

In palatal fusion, the medial-edge epithelium terminates to proliferate and enters apoptosis (Cuervo and Covarrubias, 2004; Cui et al., 2005), and the periderms covering the fusing epithelium are sloughed off (Hu et al., 2015). The intervening epithelium then needs to be degraded in order to achieve mesenchymal confluence (Gritli-Linde, 2007).

At E15.0, immunostaining for K14 revealed that epithelium seam was present sparsely at the boundary between the primary and secondary palates in the control, whereas there was partial contact but no fusion in the mutant palatal epithelium between the primary and secondary palates (Fig. 2A, B) and in the anterior-most region of the secondary palate (data not shown).

**Figure 2.**
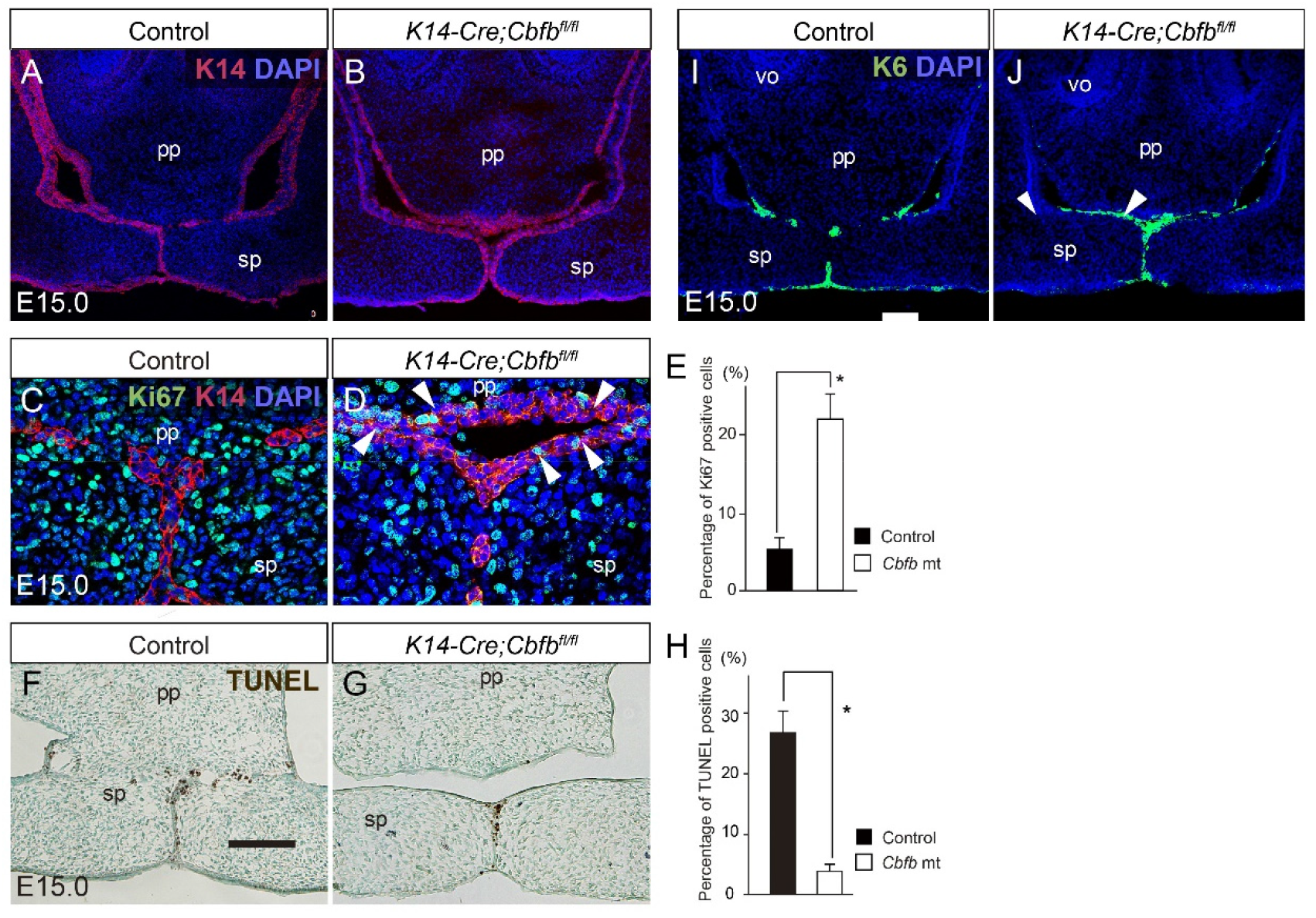
Palatal phenotypes in *Cbfb* mutant mice. (A,B) Immunostaining for K14 during the anterior palatogenesis at E15.0. In controls, K14-labeled epithelial seams were formed at the boundary between the primary and secondary palate (A). In *Cbfb* mutants, K14-labeled epithelial remnants were retained on the contacting palatal shelves (B). Scale bar: 100 μm. (C,D) Double immunohistochemical staining for Ki67 (green) and K14 (red) showed that Ki67 signals were sparse in the epithelial remnants in the wild-type palates (C), while some Ki67-positive epithelium were retained in *Cbfb* mutants (arrowheads in D). Scale bar: 50 μm. (E) There were significantly more Ki67-positive epithelial cells in the *Cbfb* mutants than in the wild-type palates. (F,G) TUNEL-positive cells were evident at the epithelial remnants localized at the boundary between the primary and the secondary palate in wild-type (F), while the epithelial remnants were less in *Cbfb* mutants (G). Scale bar: 200 μm. (H) The percentage of the TUNEL positive cells was significantly lower in the *Cbfb* mutants. (I,J) Immunohistochemical staining for K6 (green) revealed that K6-positive periderms were retained on the unfused epithelial surface of the nasal side of the secondary palate and the nasal septum. The nuclei were counterstained with DAPI (blue). The arrowhead indicates persistent periderm. The diagram shows the section positions. Scale bar: 100 μm. pp, primary palate; sp, secondary palate; ns, nasal septum.

The proliferative activity was evaluated using Ki67 staining. Double-staining for Ki67 and K14 immunoreactivity showed that Ki67-positive proliferating cells were sparse at the fused epithelium in wild-types (Fig. 2C), whereas some immunoreactivity was retained in the epithelium in *Cbfb* mutants (Fig. 2D). Ki67-positive cells were quantified from the images and we found that the percentage cells for Ki67 in the mutants was significantly higher than in the wild-type palates (Fig. 2E).

TUNEL assay showed that apoptotic signals were evident in the fused epithelium in controls (Fig. 2F), whereas far fewer signals were detected on the unfused epithelium of the *Cbfb* mutants (Fig. 2G). TUNEL-positive cells were quantified from the images and we found and TUNEL-positive cells at the fusing epithelium was significantly reduced in percentage in the mutants than in the controls (Fig. 2H).

During palatogenesis, the periderm of the secondary palate transiently covers the fusing palatal process and is sloughed before palatal fusion (Hu et al., 2015). Keratin 6 (K6) detects periderm (Richardson et al., 2014) and K6 immunoreactivity was sparsely observed in the epithelial remnants in the anterior regions of E15.0 wild-type mice (Fig. 2I). In contrast, K6-immunoreactive periderms in *Cbfb* mutants were retained on the unfused epithelial surface of the primary palate and the nasal side of the secondary palate and the nasal septum, indicating that the periderm had not been sloughed off at the anterior region of the palate by *Cbfb* deficiency (Fig. 2J).

Taken together, these findings show that *Cbfb* is essential for anterior palatal fusion and palatal fusion in *Cbfb* mutants could be due to failed disintegration of the epithelium in the anterior palate, as observed in *Runx1* mutants (Sarper et al., 2018).

### The expression of Cbfb mRNA in the developing palate

The whole-mount *in situ* hybridization showed that *Cbfb* transcripts were widely distributed along the AP axis and not specifically in the anterior regions at E14.0 (Fig. 3A,B). The distribution of the *Cbfb* mRNA expression therefore does not explain why *Cbfb* deficiency caused an anterior-specific phenotype in palatogenesis. Sliced sections revealed that *Cbfb* transcripts were present in both the palatal epithelium and mesenchymal tissue (Fig. 3C). The *Runx1* expression was intense in the fusing region of the palatal shelves and in the primary palate regions (Fig. 3D), and the *Runx2* expression was present in the fusing region of the palatal process, however, *Runx2* expression was lower in the primary palate region than the secondary palate (Fig. 3E), as previously reported (Charoenchaikorn et al., 2009). *Runx3* was also detected in the fusing region of the palatal process (Fig. 3F).

**Figure 3.**
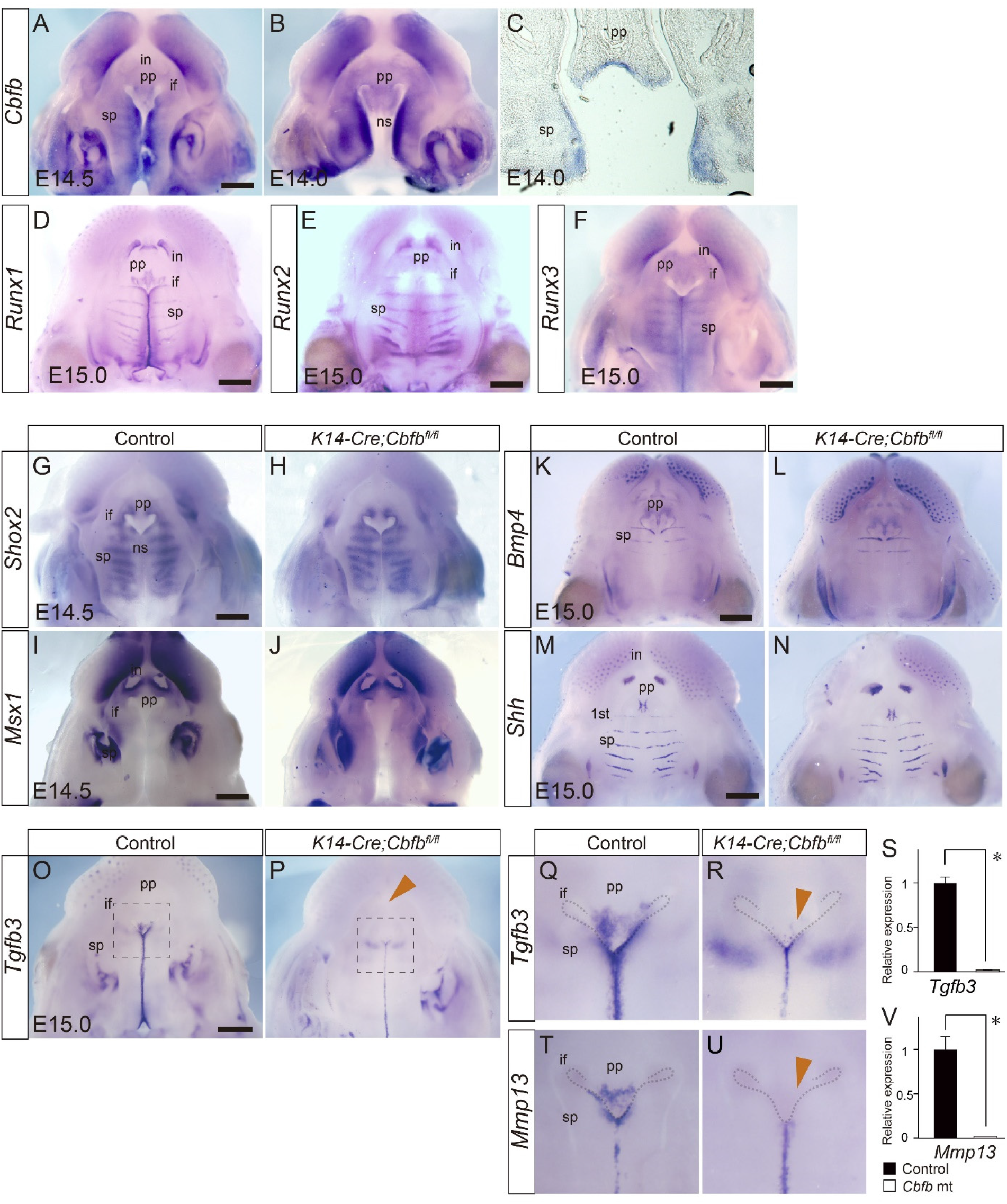
Gene expression during palatogenesis in *Cbfb* mutants. (A-F) Expression of *Cbfb, Runx1, Runx2* and *Runx3* in the developing palate of the wild-type. *Cbfb* was widely distributed along the AP axis and not specifically in the anterior regions as shown by whole-mount in situ hybridization (A,B) *Cbfb* was expressed both in the epithelium and the mesenchyme. (D-F) Whole-mount in situ hybridization of *Runx1* (D), *Runx2* (E) and *Runx3* (F) mRNA in the developing palate of wild-type mice. (G-N) Whole-mount in situ hybridization of *Shox2* (G,H), *Msx1* (I,J) and *Bmp4* (K,L) and *Shh* (M,N) mRNA in the developing palate of *Cbfb* mutant and wild-type mice. The *Shox2, Msx1, Shh* and *Bmp4* expression was not altered by *Cbfb* deficiency. (O,P) Whole-mount in situ hybridization of *Tgfb3.* The *Tgfb3* expression was markedly downregulated at the fusing epithelium at the primary palate and at the anterior-most portion of the secondary palate in *Cbfb* mutant mice. (Q,R,T,U) Higher magnification of whole-mount s images of the *Tgfb3* (inset of panel O and P) and *Mmp13.* The expression of both *Tgfb3* and *Mmp13* was markedly disturbed in *Cbfb* mutants (arrows). Scale bar: 500 μm. (S,V) qPCR analysis confirmed the remarkable downregulation of *Tgfb3* (S) and *Mmp13* (V) in *Cbfb* mutants. Scale bar: 100 μm. Error bars, *, p<0.05; pp, primary palate; sp, secondary palate; ns, nasal septum. if, incisive foramen.

### Altered mRNA expression in Cbfb mutant palate

To clarify the molecular mechanisms underlying the failed palatal fusion in *Cbfb* mutants, we evaluated the changes in several molecules that have been recognized as anterior-specific genes in palatogenesis. Whole-mount *in situ* hybridization revealed that the distribution of *Shox2, Msx2, Bmp4* or *Shh* (Baek et al., 2011; Hilliard et al., 2005; Li and Ding, 2007; Welsh and O’Brien, 2009) expression was not altered by *Cbfb* deficiency (Fig. 3G-N). However, *Tgfb3* was significantly decreased in *Cbfb* mutants in the anterior region of the palate (Fig. 3O,P). Higher magnification view demonstrated that significant decreases in *Tgfb3* signals was evident in the primary palate regions, while the *Tgfb3* expression in the secondary palate was not altered (Fig. 3Q,R). A qPCR analysis of the microdissected tissue also showed the downregulation of *Tgfb3* in the primary palate (Fig. 3S). *Mmp13* lies downstream of *Tgfb3* signaling in palatogenesis (Blavier et al., 2001). Higher magnification view of *Mmp13* expression also demonstrated that significant decreases in the signals was evident in the primary palate regions and at the anterior-most secondary palate corresponding to the 1^st^ and 2^nd^ rugae (Fig. 3T.U). qPCR analysis of microdissected tissue also confirmed marked downregulation of the expression of *Mmp13* expression in the primary palate (Fig. 3V).

These findings indicate that *Tgfb3* is one of the targets in *Cbfb* mutants, and the *Shh, Shox2* and *Msx1-Bmp4* pathways were not affected, as observed in *Runx1* mutants (Sarper et al., 2018).

### Rescue of cleft palate in Cbfb mutant mice by TGFB3

Given the critical roles of Tgfb3 in palatogenesis, downregulation of *Tgfb3* expression in Cbfb mutants might account for the failure of the palatal fusion. Therefore, we further investigated whether TGFB3 protein can rescue the cleft palate in *Cbfb* mutants. TGFB3 beads the mutant cleft rescued by 80%, while BSA treatment did not rescue it at all (4/5, Fig. 4B), indicating that *Tgfb3* is critical in the cleft palate in *Cbfb* mutants. A qPCR demonstrated that the application of TGFB3 protein resulted in upregulation of *Mmp13* expression without *Tgfb3* induction in the microdissected tissue (Fig. 4C,D). Together, these findings indicated that *Tgfb3* is a critical target in the pathogenesis of the *Cbfb* mutant cleft.

**Figure 4.**
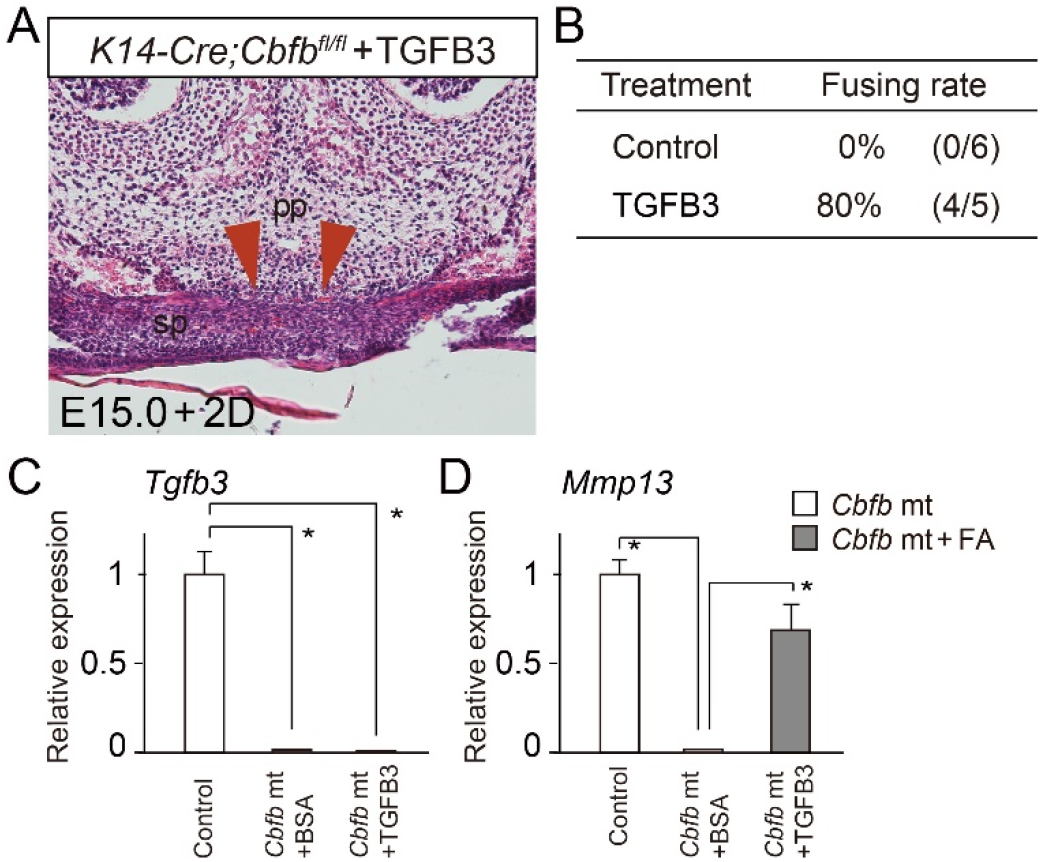
TGFB3 rescues cleft palate of *Cbfb* mutants. (A) Histological sections showed that failure of the palatal fusion in *Cbfb* mutants was partially rescued by TGFB3 protein beads in culture (Arrowheads). (B) The rescue ration of the cleft palate in *Cbfb* mutants by TGFB3 application. (C,D) qPCR analysis of the microdissected primary palate in *Cbfb* mutants demonstrated that the expression of *Tgfb3* and *Mmp13* was significantly upregulated by the folic acid application. pp, primary palate; sp, secondary palate. Error bars, *p < 0.05.

### Stat3 activity in Cbfb mutant palate

In our previous study using *Runx1* mutant mice, we demonstrated that Stat3 phosphorylation was disturbed by *Runx1* deficiency in the anterior region of the palate (Sarper et al., 2018). We therefore explored whether or not the Stat3 activity is affected during anterior palatal fusion in *Cbfb* mutants.

Immunoreactivity to Stat3 was present in the palatal epithelium, and some immunoreactivity was also observed in the mesenchyme (Fig.5A). *Cbfb* deficiency did not affect the Stat3 immunoreactivity (Fig.5B). In contrast, immunoreactivity to pStat3 was detected in the fusing or fused epithelium in wild-type (Fig. 5C), whereas pStat3 was remarkably downregulated in the primary palate in *Cbfb* mutants (Fig. 5D). A western blot analysis revealed a significant reduction in the immunoreactivity to pStat3 in the *Cbfb* mutant primary palate, while that to Stat3 was not affected (Fig. 5E).

**Figure 5.**
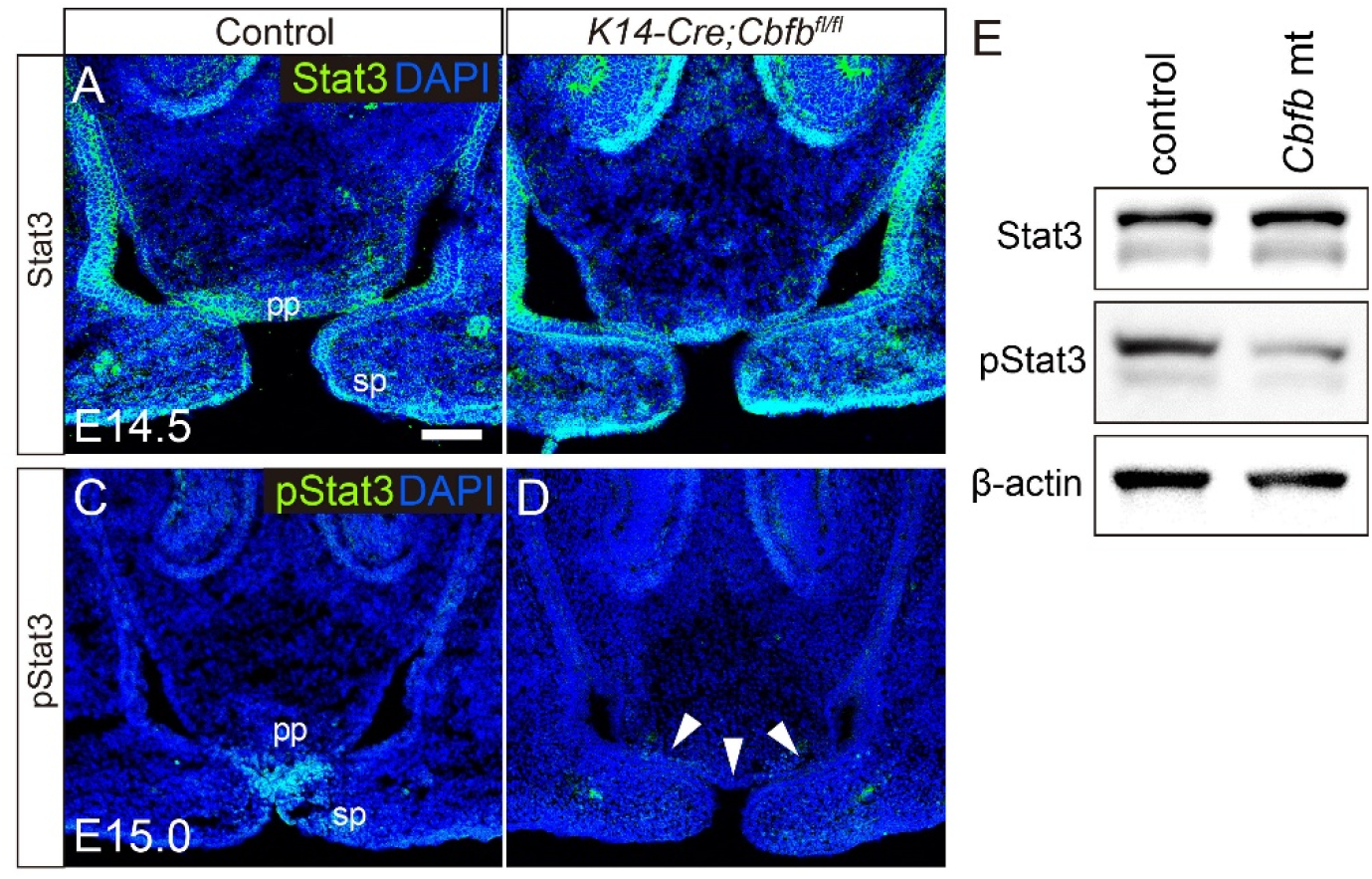
Stat3 activation in the *Cbfb* mutant palate. (A-D) Immunofluorescence analysis of Stat3 (green) and phosphorylated Stat3 (pStat3, green) in control (A, C) and *Cbfb* mutant mice (B, D). The nuclei were counterstained with DAPI (blue). pStat3 immunoreactivity was downregulated specifically at the anterior region of the palate (arrowheads in D). Scale bar: 100 μm. (E) A western blot confirmed that the pStat3 immunoreactivity was specifically downregulated in the primary palate of *Cbfb* mutants.

### Rescue of cleft palate of Cbfb mutants by folic acid

We then attempted to rescue the mutant cleft palate using folic acid application. A recent study showed that folic acid and folate activate STAT3 pathway (Hansen et al., 2015; Wei et al., 2017). We therefore investigated whether or not folic acid application could rescue the anterior cleft palate of *Cbfb* mutants.

After 48 h application of folic acid, histological observation confirmed the partial achievement of mesenchymal continuity by folic acid application in the mutant palatal explants (Fig. 6A). Folic acid application rescued the failed palatal fusion with a success rate of 67% (4/6, Fig. 6B). A western blot showed that folic acid activated pStat3 immunoreactivity, while the Stat3 was not altered in the dissected mutant primary palate (Fig. 6C). qPCR of the micro-dissected primary palate revealed that the expression of *Tgfb3* and *Mmp13* was upregulated by folic acid application (Fig. 6D,E).

**Figure 6.**
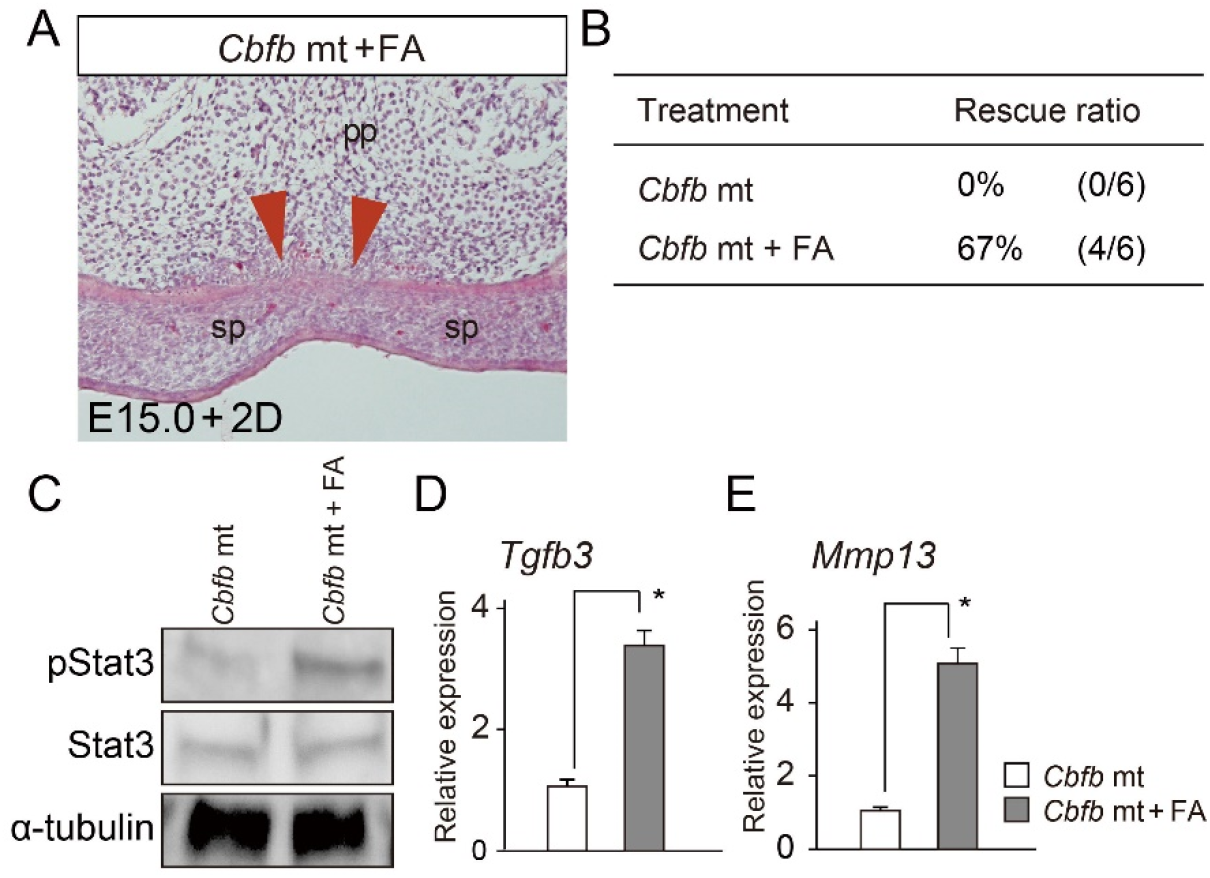
Folic acid application rescues cleft palate of *Cbfb* mutants. (A) Histological sections showed that failure of the palatal fusion in *Cbfb* mutants was partially rescued by folic acid application in culture (Arrowheads). (B) The rescue ration of the cleft palate in *Cbfb* mutants using folic acid. (C) A western blot analysis confirmed that the pStat3 immunoreactivity was upregulated by folic acid application (FA) in the primary palate of *Cbfb* mutants. (D,E) qPCR demonstrated folic acid application (FA) significantly upregulated in the palatal tissues of *Cbfb* mutants *in vitro.* pp, primary palate; sp, secondary palate. Error bars, *p < 0.05.

## Discussion

This present study provides the first genetic evidence of Cbfb being necessary for palatogenesis using conditional *Cbfb* null mutant mice. *Cbfb* deficiency resulted in anterior cleft between the primary and secondary palate and led to the failed disintegration of the contacting palatal epithelium as observed in *Runx1* mutants (Sarper et al., 2018). Cbfb forms a heterodimer with *Runx* genes. In hematopoietic development, the functional loss of either *Runx1* or *Cbfb* completely disturbed the function in hematopoietic cells, indicating that Cbfb act as an obligate cofactor for the Runx function (Chen et al., 2011; Chen et al., 2009; Gau et al., 2017). In contrast, *Cbfb* deficiency does not completely disturb the Runx2-dependent bone and cartilage formation (Yoshida et al., 2002), suggesting that Runx2 can regulate skeletogenesis to a limited degree even in the absence of *Cbfb* (Gau et al., 2017), and Cbfb acts as a dispensable modulator of Runx activity in skeletogenesis (Gau et al., 2017). Given the similarities in the anterior cleft palate observed after the loss of function of *Cbfb* or *Runx1,* Cbfb appears to serve as an obligate cofactor, rather than a modulator in Runx1/Cbfb signaling during palatogenesis.

Our findings also provide the additional evidence that Runx signaling is important in the anterior palatogenesis and that Tgfb3 is a critical downstream target. As observed in *Runx1* mutants (Sarper et al., 2018), *Tgfb3* expression was specifically downregulated in the *Cbfb* mutants and conversely, TGFB3 protein beads rescued the failed palatal fusion in the mutant. Indeed, epithelial-specific depletion of *Tgfb3, Tgfbr1 (Alk5),* or *Tgfbr2* results in anterior-specific palatal cleft (Dudas et al., 2006; Lane et al., 2015; Xu et al., 2006). On the other hand, pharmaceutical Stat3 inhibitor also disturbs the anterior palatal fusion with marked downregulation of *Tgfb3* expression (Sarper et al., 2018) and we found that Stat3 phosphorylation was also disturbed in *Cbfb* mutants. Given that the obligatory roles of Cbfb in Runx1 signaling, the downregulation of *Tgfb3* in the primary palate may account for the anterior-specific clefting in *Cbfb* mutants, as observed in *Runx1* mutants. In addition, these findings are the additional evidences that support the essential roles of Runx1/Cbfb-Stat3-Tgfb3 signaling axis in anterior palatogenesis (Fig. 7A-C).

**Figure 7.**
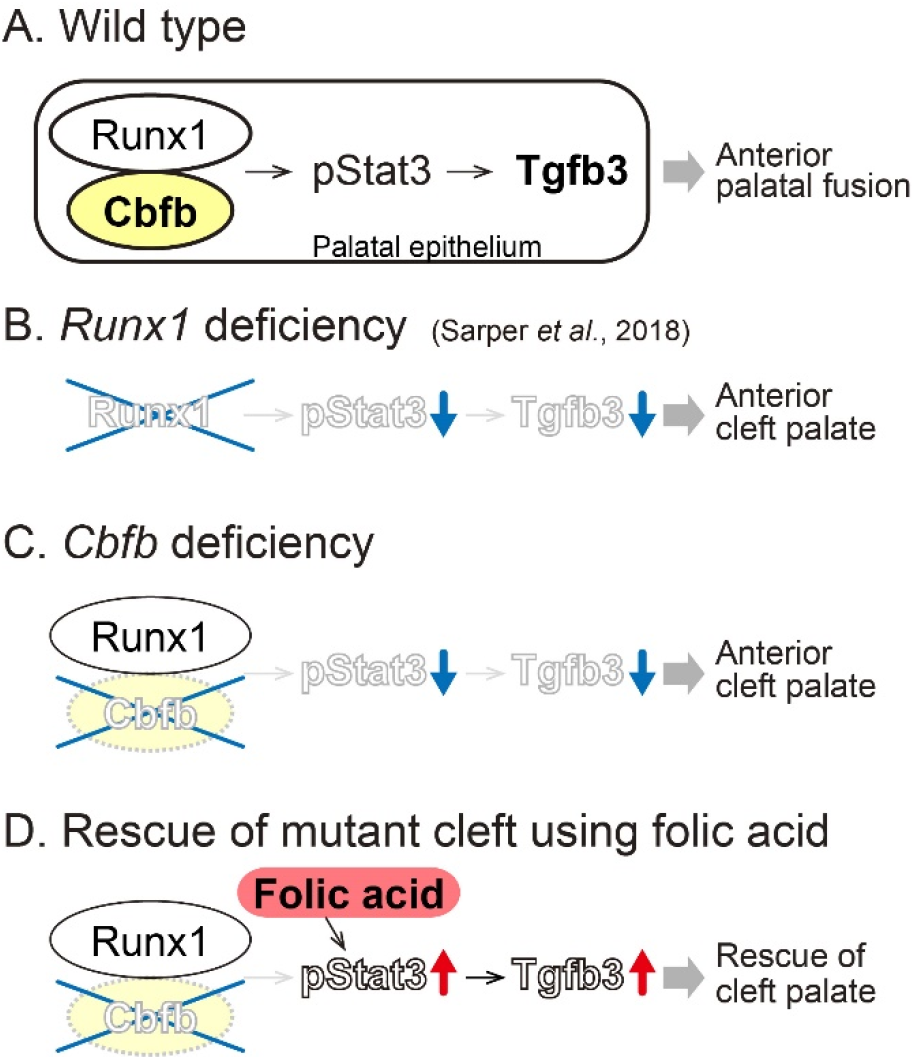
A schema of the key findings. Runx1/Cbfb-Stat3-Tgfb3 signaling regulates the fusion of the anterior palate. (A) In the fusing palatal epithelium of the wild-type palate, Runx1/Cbfb is involved in regulation of Stat3 phosphorylation, which further regulates the *Tgfb3* in the anterior region of the palate. (B) *Runx1* mutants exhibit anterior clefting and *Tgfb3* expression is remarkably disturbed in the primary palate with downregulation of Stat3 phosphorylation, as shown previously (Sarper et al., 2018). (C) *Cbfb* mutant mice also display an anterior cleft palate with the downregulate *Tgfb3* expression and the suppressed Stat3 phosphorylation. (D) The anterior cleft palate in *Cbfb* mutants is rescued by pharmaceutical application of folic acid that activates suppressed Stat3 phosphorylation and *Tgfb3* expression.

One of the more striking findings is that the folic acid application rescued the cleft palate in *Cbfb* mutants. In humans, maternal folic acid supplementation has been proven an effective intervention for reducing the risk of non-syndromic cleft palate (Millacura et al., 2017; Wehby and Murray, 2010). However, the mechanism by which folic acid prevents structural anomalies in the fetus is still unknown (Obican et al., 2010). A recent study showed that folic acid can activate Stat3 (Hansen et al., 2015; Wei et al., 2017). In the present study, phosphorylation of Stat3 was activated by folic acid application in the dissected palatal tissue in culture. Conversely, a Stat3 inhibitor impairs anterior palatal fusion between the primary and secondary palates and disturbed the expression of *Tgfb3 in vitro* (Sarper et al., 2018). Taken together, these findings show that folic acid rescued the cleft palate of *Cbfb* mutants, presumably through the activation of Stat3. Furthermore, the rescue of the mutant cleft palate using folic acid may elucidate potential therapeutic targets by Stat3 modification for the prevention and pharmaceutical intervention of cleft palate (Fig. 7D).

In conclusion, the present study demonstrated that Cbfb is essential for anterior palatogenesis as an obligatory cofactor of Runx1/Cbfb signaling (Fig. 7A). In addition, we also demonstrate the rescue of mutant cleft palate via pharmaceutical folic acid application, at least in part, by activating Stat3 phosphorylation in the Runx/Cbfb-Tgfb3 signaling axis during palatogenesis (Fig.7D).

## Materials and Methods

### Animals

*Cbfb^-/-^* mice are early lethal due to hemorrhaging between E11.5 and E13.5, when the palatal development is not yet initiated (Sasaki et al., 1996). To assess the role of Cbfb in the oral epithelium, we use epithelial-specific knock-out mice created through the *Cre/loxP* system (*K14-Cre/Cbfb^fl/fl^),* as described in a previous study (Kurosaka et al., 2011). We used their littermates that did not carry the *K14-Cre/Cbfb^fl/fl^* genotype as controls.

### Assessment of palatal fusion and a histological analysis

The palatal phenotypes were first evaluated with a dissecting microscope. For histology, dissected samples were fixed in 4% paraformaldehyde at 4 °C overnight. The samples were then dehydrated, embedded in paraffin, serially sectioned at 7 μm, and stained with hematoxylin and eosin. For cryosections, the samples were dehydrated in 15% and 30% sucrose in DEPC-treated PBS overnight at 4 °C, embedded in Tissue-Tek (OCT compound, Sakura). The tissue samples were sectioned into 10 μm slice.

### Immunohistochemistry

Immunofluorescence staining was performed using polyclonal rabbit-anti-Ki67 (1:400, ab15580, Abcam), polyclonal rabbit anti-K6 (1:200, #4543, 905701, Biolegend), monoclonal anti-K14 (1:200, ab7880, Abcam), monoclonal rabbit anti-phospho-Stat3 (pStat3, 1:200, #9145, Cell Signaling Technology), monoclonal rabbit anti-Stat3 (1:200, #9139, Cell Signaling Technology) overnight at 4°C. Alexa488-conjugated goat-anti-rabbit IgG (1:400, A21206, Molecular Probes) or Alexa546-conjugated goat-anti-mouse IgG (1:400, A11003, Molecular Probes) was used as secondary antibody. DAPI (1:500, Dojindo) was used for nuclear staining and the sections were mounted with fluorescent mounting medium (Dako). At least three embryos of each genotype were used for each analysis.

The percentage of proliferating cells at the fusing or contacting epithelium between the primary and the secondary palate was determined by counting Ki67-positive cells as a percentage of the total number of cells, as determined by DAPI staining.

### Laser microdissection

The dissected heads were freshly embedded in OCT compound (Tissue Tek, Sakura) and frozen immediately. Then, tissues were serially sectioned at a thickness of 25 μm on a cryostat (Leica CM 1950). Theater, the sections were mounted on a film-coated slide. From the anterior palate at E15.0, 12-14 serial sections were obtained on total and stained with Cresyl violet. Palatal tissues at the border between the primary and the secondary palate were dissected from the sample sections using a manual laser-capture microdissection system (LMD6500, Leica) and collected into tubes.

### RNA Extraction and a real-time RT-PCR Analysis

Total RNA was extracted from the laser-microdissected tissues or dissected tissues using IsogenII (Nippon Gene, Toyama, Japan) according to the manufacturer’s protocol, then reverse transcribed to cDNA using an oligo (dT) with reverse transcriptase (Takara, Osaka, Japan). For real-time RT-PCR, the cDNA was amplified with TaqDNA Polymerase (Toyobo Sybr Green Plus, Osaka, Japan) using a light cycler (Roche). *Gapdh* was used as a housekeeping gene. Primer sequences are shown previously (Sarper et al., 2018). At least three embryos of each genotype were used for each analysis.

### Whole-mount in situ hybridization

Whole-mount in situ hybridization was performed using fixed E14.0, E14.5 and E15.0 palates. The digoxigenin-labeled RNA probes were prepared using a DIG RNA labeling kit according to the manufacturer’s protocol (Roche) using each cDNA clone as the template. The probes were synthesized from fragments of *Cbfb, Runx1, Runx2, Runx3, Shox2, Msx1, Shh, Bmp4, Tgfb3,* and *Mmp13* (Allen Institute for Brain Science) and amplified with T7 and SP6 adaptor primers through PCR, as described previously (Sarper et al., 2018). After hybridization, the signals were visualized according to their immunoreactivity with anti-digoxigenin alkaline phosphatase-conjugated Fab fragments (Roche). At least three embryos of each genotype were used for each analysis.

### TUNEL staining

To detect apoptotic cells, the TUNEL assay was performed according to the manufacturer’s instructions (ApopTag; Chemicon). Frozen sections (10 μm) were prepared and the stained sections were counterstained with methyl green. At least three embryos of each genotype were used for each analysis.

The percentage of apoptotic cells along the contacting or fused epithelium between the primary and the secondary palate was determined by TUNEL-positive cells as a percentage of the total number of cells, as determined by methyl green staining.

### Rescue of the mutant clef palate using TGFB3 protein or folic acid

The dissected palate of the E15.0 mutants was cultured on a Nuclepore filter (Whatman, Middlesex, UK) in Trowell type organ culture in chemically defined medium (BGJb: gibco /life technologies). Affi-Gel beads (Bio-Rad) were incubated in TGFB3 (100 ng/μl, R&D Systems) and placed on the primary palate of the *Cbfb* mutant explants, as described previously (Sarper et al., 2018). Bovine serum albumin (BSA) was used for the control beads. Fusion of the palatal process was evaluated histologically. The anterior portion of the palates was also dissected under the microscope and total RNA was extracted from these samples for pPCR analysis.

To evaluate the possible rescue of cleft palate in Cbfb mutants by folic acid application, the palatal explants were cultured for 48 h in BGJb (Gibco) culture medium containing folic acid (N^5^-formyl-5,6,7,8-tetrahydropteroyl-L-glutamic acid) (Sigma) at a final concertation of 100 μg/ml. After culture, the *in vitro* explants were fixed at each stage in 4% paraformaldehyde overnight and then processed for histological observation.

### Western blot analysis

For western blotting, the primary palate of *Cbfb* mutants was dissected and then cut in half. Each half of the explants was cultured with or without folic acid for 48 hours.

The dissected samples were lysed with RIPA buffer (nacalai tesque) supplemented with protease and phosphatase inhibitors (nacalai tesque). The lysates were centrifuged and the supernatant was heated in denaturing Laemmli buffer (Bio-rad Laboratories). Proteins were separated by SDS-PAGE and transferred to polyvinylidene difluoride membranes (Bio-rad Laboratories).

The membranes were incubated with anti-Stat3 (1:1000, #9139, Cell Signaling Technology), anti-pStat3 (1:1000, #9145, Cell Signaling Technology), anti-β-actin (1:2000, Sigma) or anti-α-tubulin (1:1000, Invitrogen). The bound antibodies were detected with HRP-linked antibody (1:1,000, Cell Signaling Technology) and an ECL detection kit (Bio-rad Laboratories).

### Statistical analyses

Quantitative variables in the two groups were compared using the Mann-Whitney *U* test. Differences among the three groups were determined using the analysis of variance (ANOVA) test, and significant effects indicated by the ANOVA were further analyzed with post hoc Bonferroni correction. *P* values < 0.05 were considered significant. Significance was determined using the statistical analysis software program JMP, version 5 (SAS Institute Inc.)

### Study approval

All of the animal experiments were performed in strict accordance with the guidelines of the Animal Care and Use Committee of the Osaka University Graduate School of Dentistry, Osaka, Japan. The protocol was approved by the Committee on the Ethics of Animal Experiments of Osaka University Graduate School of Dentistry. Mice were housed in the animal facility at the Department of Dentistry, Osaka University. Welfare guidelines and procedures were performed with the approval of the Osaka University Graduate School of Dentistry Animal Committee.

## Acknowledgments

We thank Ms. Yuriko Nogami for the excellent care and maintenance of our mouse colony and her valuable assistance in the histological, molecular and protein work.

## Author contributions statement

T.Y. designed the study. S.E.S., T.I., H.K., H.O.M., Y.M. and T.S. performed and/or analyzed experiments. I.T. and K.K. provided experimental reagents and participated in the discussions. T.Y. and S.E.S. wrote the manuscript with input from all authors. All authors read and approved the final manuscript.

